# Missing-value imputation and *in-silico* region detection for spatially resolved transcriptomics

**DOI:** 10.1101/2021.05.14.443446

**Authors:** Linhua Wang, Zhandong Liu

## Abstract

We are pleased to introduce a first-of-its-kind algorithm that combines *in-silico* region detection and spatial gene-expression imputation. Spatial transcriptomics by 10X Visium (ST) is a new technology used to dissect gene and cell spatial organization. Analyzing this new type of data has two main challenges: automatically annotating the major tissue regions and excessive zero values of gene-expression due to high dropout rates. We developed a computational tool—MIST—that addresses both challenges by automatically identifying tissue regions and estimating missing gene-expression values for individual tissue regions. We validated MIST detected regions across multiple datasets using manual annotation on the histological staining images as references. We also demonstrated that MIST can accurately recover ST’s missing values through hold-out experiments. Furthermore, we showed that MIST could identify subtle intra-tissue heterogeneity and recover spatial gene-gene interaction signals. We therefore strongly encourage using MIST prior to downstream ST analysis because it provides unbiased region annotations and enables accurately de-noised spatial gene-expression profiles.

## Introduction

To understand the biological mechanisms underlying diseases, it is essential to delineate cell and gene spatial organizations; the scientific community has therefore invested significant time and effort in detecting positional gene-expression^1,2^. Of all the spatial gene-expression profiling techniques that have been developed, 10X Visium Spatial Transcriptomics (ST) gained its most popularity due to its whole-genome scalability and cost-efficiency^2,3^. So far, ST has been used to study several tissues’ spatial gene-expression, including breast cancer, prostate tumors, melanoma tumors, human and mouse brains, and more^3–8^.

Despite its success and popularity in profiling positional gene-expression, ST data analysis yields a large number of zero-values due to technical dropouts^3^. When these dropout events exist in spatially resolved transcriptomics, it prevents accurate cluster detection and downstream analysis and drastically reduces the single-to-noise ratio in the data. Accurately estimating dropped-out expression values in ST data is therefore essential to rescuing such signals and facilitating more accurate spatial-gene-expression-pattern detection.

While we lack imputation methods specifically designed to impute missing values in ST, plenty of imputation methods have been developed for single-cell RNA sequencing (scRNA-seq) data^9–14^. Many of these methods— like knnSmoothing, MAGIC, and PRIME—rely on defining a cell affinity matrix or even forming small local networks purely based on the molecular profile^9–11^. They cannot, however, interpret ST's spatial connectivity information to determine affinity inference.

An intuitive way of imputing ST using spatial information is to use spatially adjacent neighbors to estimate missing values. However, using adjacent neighbors introduces errors regarding spots on functionally different tissue regions’ boundaries. Detecting the tissue boundaries is vital to avoiding spatial-information misusage when imputing ST data and allows ST to be grouped into functionally similar regions. Imputing within such functional and spatial subregions therefore avoids errors at tissue boundaries.

When assigning coordinates to the transcriptomes sequenced from the tissue, the tissue boundaries will not be clear in the ST data; this poses another challenge inherent in many ST data analyses. We routinely use the naked eye to perform manual annotation and predict tissue regions, which can cause false classifications. Automatically detecting tissue boundaries and regions from the molecular profile and spatial information is therefore essential to facilitate region-specific imputation and many other downstream analyses.

Our new tool – Missing-value Imputation for Spatially resolved Transcriptomics (MIST), first detects spatial and functional boundaries from the observed spatial gene-expression matrix and then uses the rank minimization algorithm to impute each detected tissue region (Fig. 1). We assume that each detected tissue region has a low-rank representation because of the small number of cell types within them. MIST can utilize the heterogeneity within each region to perform more accurate imputation. Moreover, by introducing noises into each major region and using ensemble learning to stabilize the outcomes from multiple repeats, MIST will more robustly estimate the missing values.

**Figure 1.**
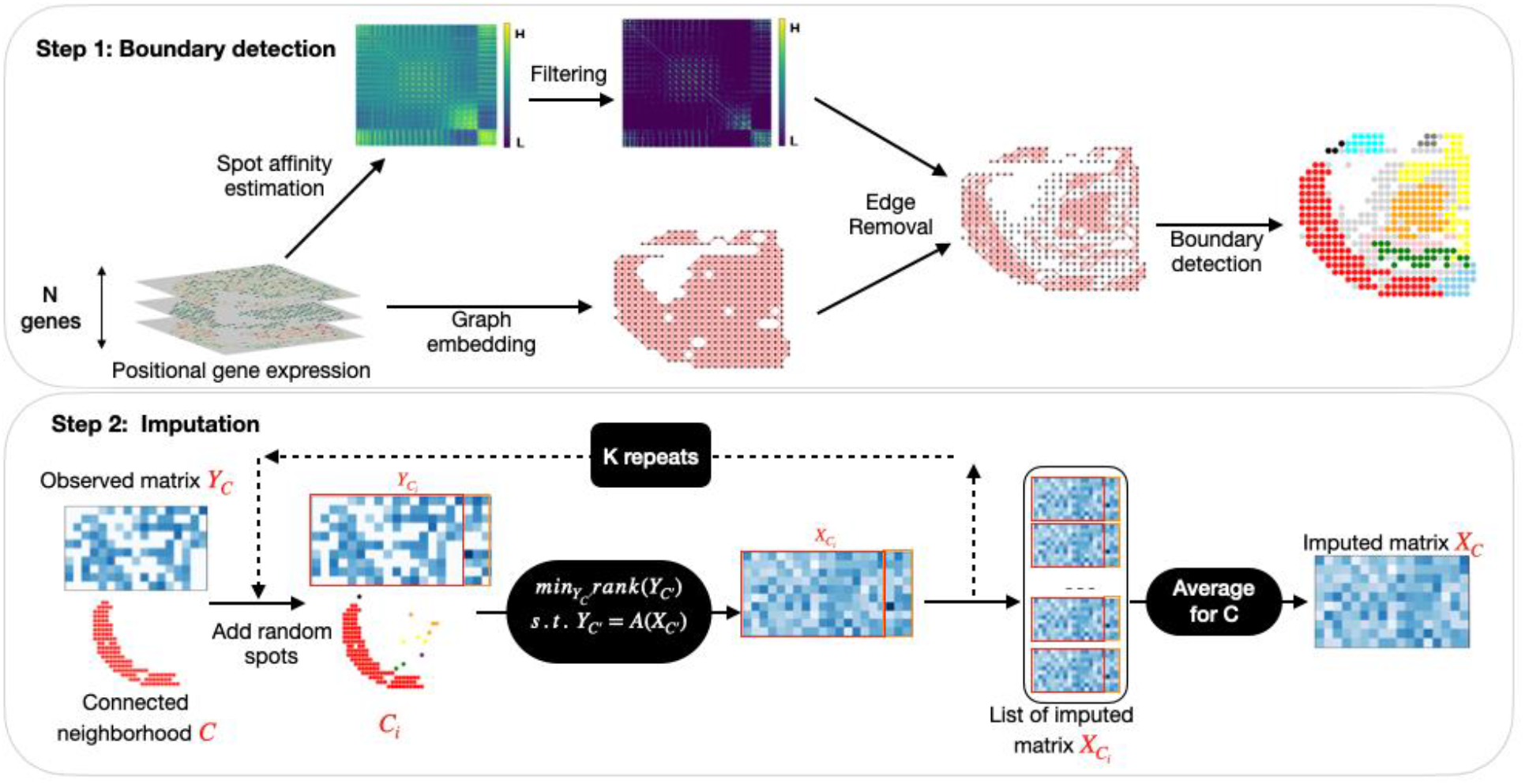
The MIST pipeline. In step 1, sub-region boundaries are detected by extracting local connected components (LCN) through graph embedding and edge filtering using molecular similarity. In step 2, missing values for each LCN are estimated using the average of multiple runs of rank minimization algorithm with the augmented region.

## Results

### The MIST Algorithm

MIST uses molecular features and spatial connectivity to identify functional spatial regions, and then uses the average of multiple rank-minimization-approach runs to impute missing values for each detected region with augmented spots (Fig. 1).

MIST first detects functional spatial regions by extracting locally connected neighbors (LCNs). It uses the CPM-normalized high-dimensional gene-expression matrix as input and embeds the spots in a 2D graph with edges that connect only adjacent spots. It then uses Principal Component Analysis (PCA) to project the data into low-dimension space and remove the noise^15^. The Pearson Correlation Coefficient (PCC) between the top principal components captures the weight of each edge that connects two spots. Edges with low weights are removed and the filtering threshold is automatically selected using a high-coverage landmark gene set.

After detecting the LCNs, MIST estimates the missing values in each detected LCN by averaging the outcomes from multiple rank-minimization-algorithm runs with augmented random spots (Algorithm 1). We first assume that each LCN’s gene-expression matrix should have a low rank due to the small number of cell types. By augmenting each LCN with random spots and averaging the outcomes after multiple runs, we added signals from outside the LCN to make our imputed results more robust to noise.

### MIST detected functional regions within tissues

While higher similarity thresholds on the edges will find strongly correlated Local Connected Neighbors (LCNs), they also lead to a significant number of isolated spots with no connected edges (Fig. 2c). But, while small thresholds will include most spots, they will lose the specificity that allows us to accurately detect tissue regions (Fig. 2c). Balancing LCN-capture-proportion and LCN-detection-accuracy is, therefore, crucial to successful region detection and LCN-based imputation.

**Figure 2.**
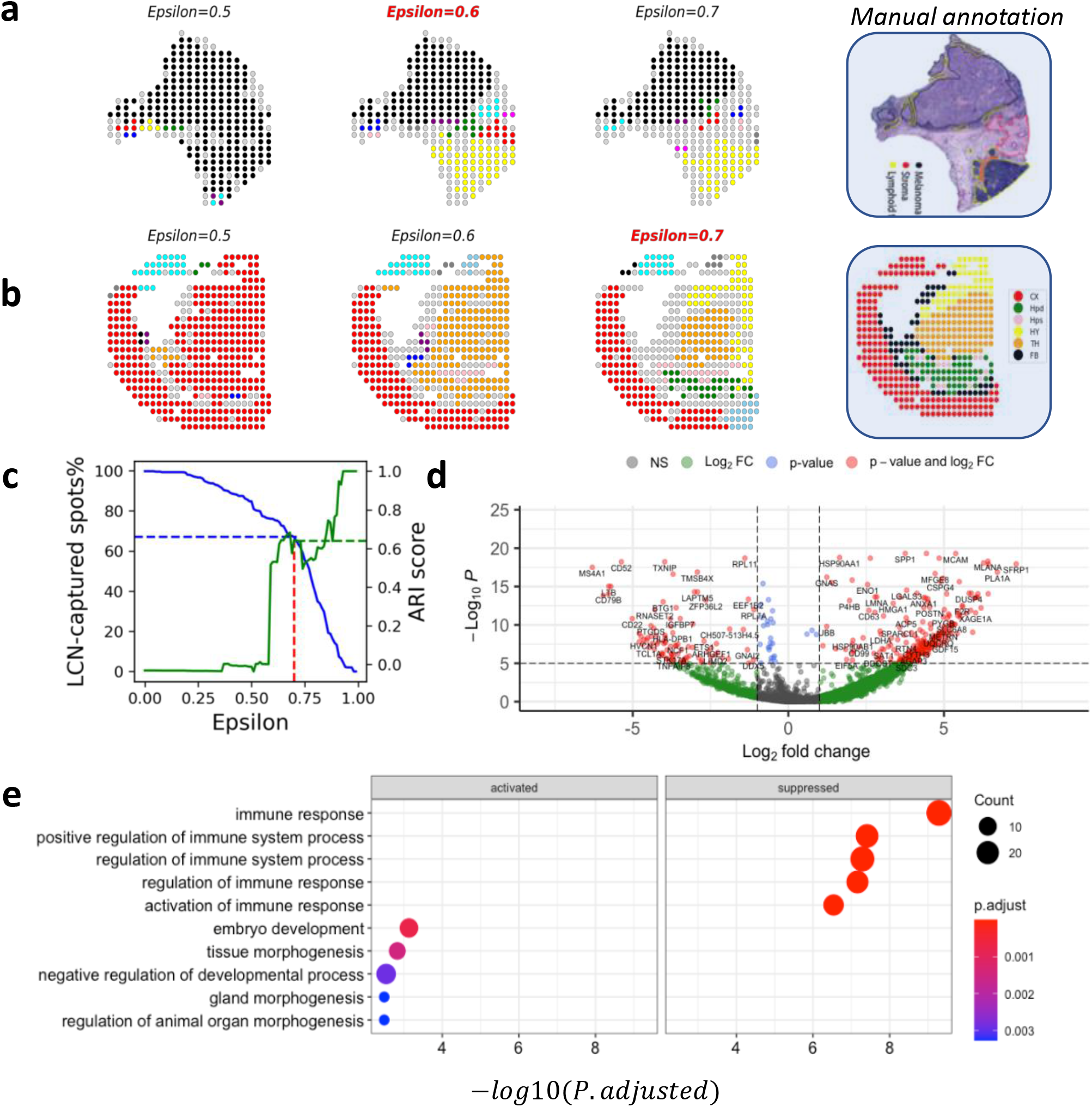
MIST detected regions agree with manual annotations. **a,** Melanoma tissue regions detected using different epsilon values. Column 2 with epsilon of 0.6 was selected by MIST. Column 4 is the pathological annotation on the aligned histological staining image. **b,** Mouse brain tissue regions detected using different epsilon values. Column 3 with epsilon of 0.7 was selected by MIST. Column 4 is the spot-level manual annotation from the original study. **c,** Percentage of spots for every component in the mouse brain sample where light gray color represents isolated spots. **d,** Percentage of spots (left y-axis, blue curve) and adjusted rand index (ARI, right y-axis, green curve) as functions of edge-filtering parameter epsilon. **e**, Volcano plot of differential expressed genes in tumor region contrasted to lymphoid region of the Melanoma sample.

We demonstrated that MIST strikes this balance automatically and accurately. We used Adjusted Rand Index (ARI) to evaluate the consistency between the regions detected under different thresholds and the human experts’ annotations. Using the Mouse WT brain sample, we validated that the threshold that MIST automatically selected found a balance point that maximize the ARI while minimizing the proportion of isolated spots (Fig. 2c). Using the selected threshold, the mouse brain regions MIST detected agreed with the anatomical regions (Fig. 2b) with an adjusted rand index (ARI) value of 0.64.

Moreover, using the optimized threshold, MIST detected the pathological regions in the Melanoma Tumor (Fig 2a, black, yellow, and red) that match the manual pathological annotation of Tumor, Lymphoid and Stroma regions on the Hematoxylin and Eosin stained image^6^.

We further validated the regions detected in the Melanoma tumor by identifying differentially expressed genes between the detected tumor region and lymphoid region. Specifically, we used the Wilcoxon rank-sum test to select significantly activated genes in tumor regions relative to lymphoid regions and vice versa. By selecting genes with a log2 fold-change (LFC) greater than 0.58 and an adjusted P-value less than 1E-5, we found 143 and 49 marker genes for the tumor and lymphoid regions, respectively (Fig. 2d). We observed that some well-known melanoma marker genes, including *MLANA^16^*, MCAM^17^, *SPP1^18^* and *HSP90AA1*^19^, topped the list of the melanoma-activated genes that we identified, which demonstrates our detected region’s accuracy. We then performed gene set enrichment analysis using R package *clusterProfiler*^20^ and observed that the immune response gene ontology term was significantly enriched (FDR = 9E-8, Fig. 2e) for the detected region that matches manual lymphoid-region annotation. Taken together, the region automatically detected from the Melanoma ST reflected the hidden functional regions within the tissue.

### MIST accurately recovers hold-out values across multiple data sets

To assess MIST’s accuracy when estimating missing values, we performed random hold-out experiments in which we withheld a random set of the observed non-zero values and used these as ground truth to evaluate the models’ performances. By withholding some of the observed values, we simulated cases in which non-zero expression values have dropped out. We compared MIST with state-of-the-art scRNA-seq methods, including MAGIC, knn-smoothing, SAVER, and McImpute, and a baseline k-nearest neighbor method (spKNN) that estimates missing values by averaging spatially adjacent neighbors.

We tested performance on four datasets from samples from a Mouse WT brain, Mouse AD brain, Melanoma tumor, and Prostate tumor. For each dataset, we selected genes with 50% non-zero values across all spots to generate hold-out test data sets. For each gene, we partitioned the non-zero expression values into five non-overlapping sets. Then, we iteratively held out one-fold of the non-zero values and assessed MIST and other imputation methods' accuracy in recovering the held-out values.

We used RMSE and PCC to evaluate imputation accuracy. Better imputation methods are expected to have lower RMSE and higher PCC scores.

We found that MIST consistently outperformed other methods across all datasets with higher PCC and lower RMSE scores during hold-out value evaluation (Fig. 3a, b). For the spots that are members of major components, MIST had an average RMSE improvement (decreasing values) of 12% (p-value=3E-4) compared with mcImpute, and 14% (p-value=1E-14) and 35% (p-value=7E-10) compared with the baseline spKNN algorithm. Moreover, SpImpute’s mean PCC is also 8% (p-value=9E-6), 4% (p-value=2E-4), and 16% (p-value=6E-11) higher than McImpute, MAGIC and spKNN respectively. Knn-smoothing and SAVER consistently performed significantly worse than the other four methods (Ext. Fig. 3.1, 3.2).

**Figure 3.**
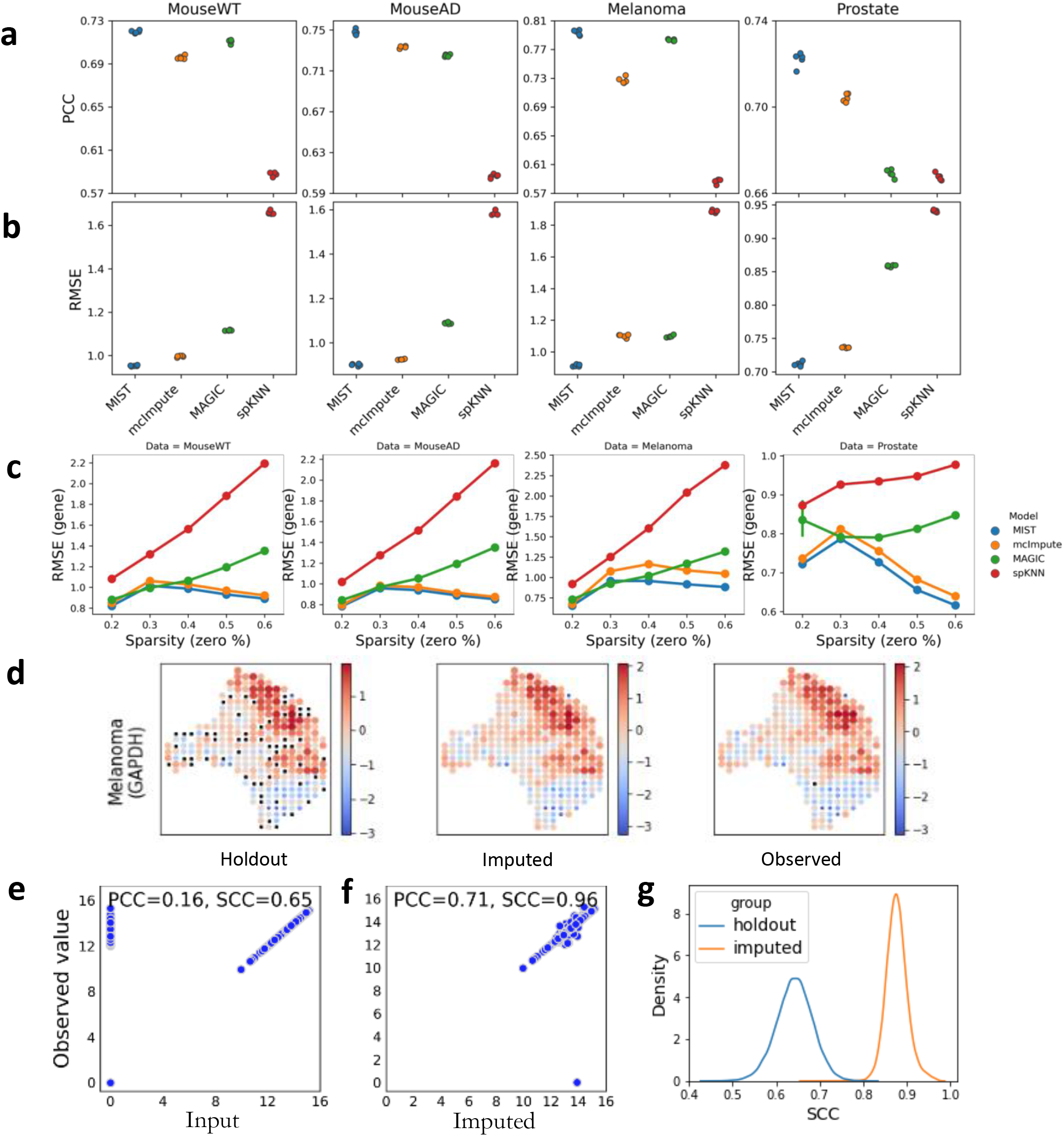
MIST’s outperforms other imputation methods in holdout experiments. **a-b**, Holdout experiment performance across multiple data sets using metrics **a**, PCC and **b**, RMSE. Each column is an individual data set. Points in the same color indicate the 5 non-overlap fold of tests for each model. **c**, Gene-level performance of each model represented by the RMSE (y-axis) as a function of zero-value percentage of the gene expression values (sparsity, x-axis). **d**, Expression patterns recovered for gene GAPDH in the Melanoma sample. From left to right shows the spatial pattern of the input to MIST with holdouts, the imputed gene expression pattern and the original observed gene expression pattern. Color and size indicate relative gene expression abundance. **e**, the correlation between holdout input and observed GAPDH expression values. **f,**the correlation between MIST imputed and observed GAPDH expression values. **g**, Comparison of distribution of the correlation with non-zero original observed values for all genes. Blue: holdout input vs. original observed. Orange: imputed vs. original observed.

We further stratified the evaluation at a per-gene level grouped by the zero-value proportion (sparsity). While MAGIC and spKNN’s RMSE monotonically increased with sparsity level, MIST’s performance was not vulnerable to gene sparsity (Fig. 3c). Though McImpute’s performance was also not influenced by gene sparsity, MIST has a superior performance than McImpute at every gene sparsity level (Fig. 3c).

We then visualized sample genes’ spatial patterns to demonstrate that MIST recovers the gene-expression patterns using the held-out data as input. Specifically, we showed that MIST can faithfully recover the gene-expression spatial pattern for gene *GAPDH* after imputation using the Melanoma tissue sample (Fig. 3d). With the hold-out input, MIST accurately estimated the original expression values by increasing the Spearman’s Correlation Coefficient (SCC) from 0.65 to 0.96 and the PCC from 0.16 to 0.71 (Fig. 3e). When evaluating all genes across the four tested data sets, after imputation, the median SCC had significantly improved from 0.64 to 0.88 (p-value=0, Fig 3f).

### MIST discovered intra-cortex heterogeneity within an Alzheimer’s Disease (AD) mouse brain

Next, we evaluated MIST by using Uniform Manifold Approximation and Projection (UMAP)^22^ to see the clustering structures. First, we performed UMAP on Mouse WT unimputed and imputed data to observe the clusters. After imputation, the heterogeneity within the brain enhanced within the cortex, somatic layer of hippocampus and thalamus, forming individual clusters (Fig. 4a).

**Figure 4.**
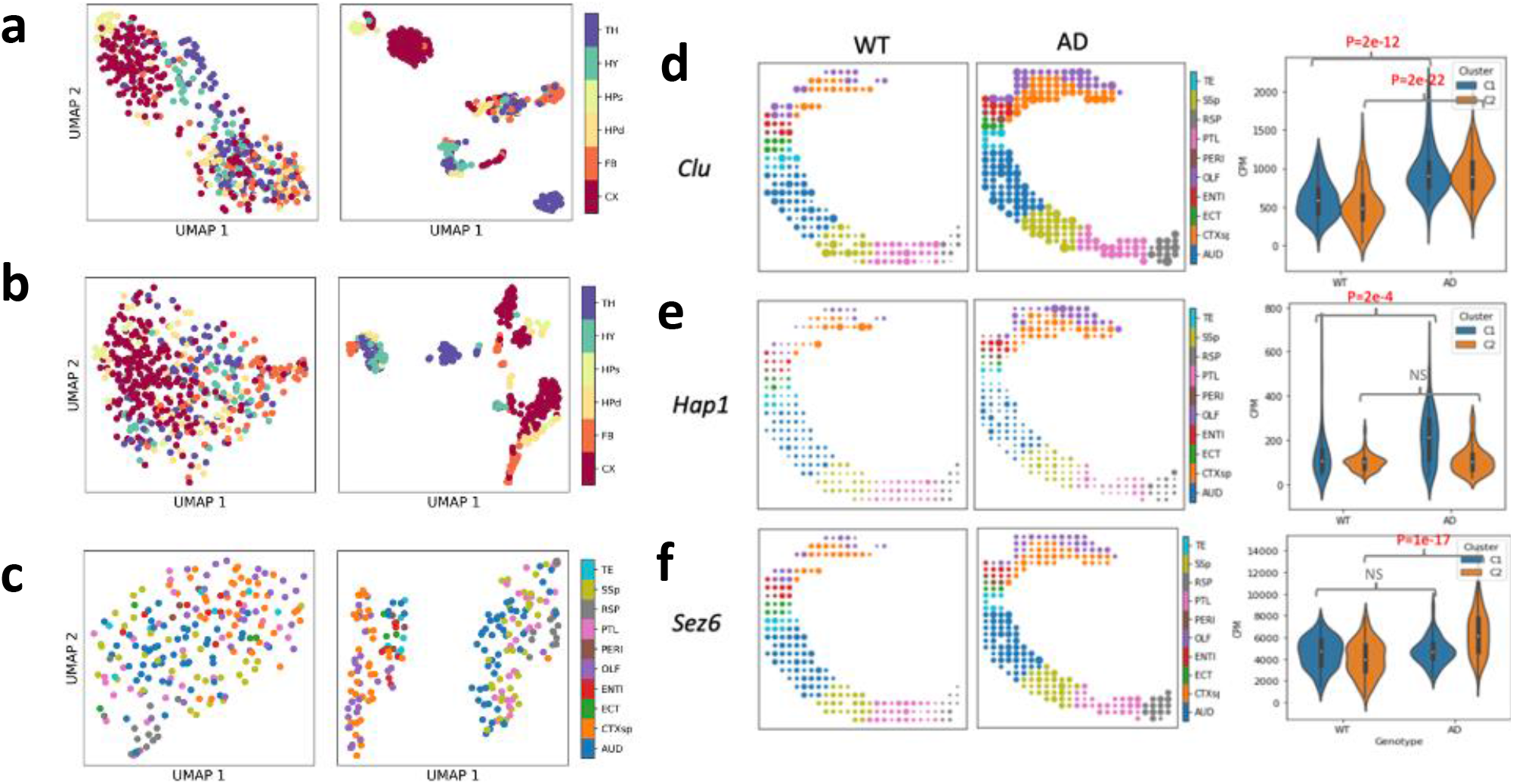
MIST identified intra-cortex heterogeneity within Alzheimer’s Disease (AD) mouse brain. **a,** UMAP of Mouse WT brain using original observed ST data (left) and MIST-imputed ST data (right). **b,** UMAP of Mouse AD brain using original observed ST data (left) and MIST-imputed ST data (right). **c,** UMAP of Mouse AD cortex region using original observed ST data (left) and MIST-imputed ST data (right). **d-f,** Examples of AD up-regulated genes that are activated in both clusters (**d**), only significantly upregulated in Cluster 1 (**e**, CTXsp, OLF, ENTI, TE, ECT and PERI) and Cluster 2 (**f**, AUD, PTL, RSP, SSp). Left, spatial expression pattern in WT mouse brain cortex; middle, spatial expression pattern in AD mouse brain cortex; right, boxplot of CPM grouped by genotype and spatial clusters.

When evaluating the Mouse AD brain using UMAP, we observed that the cortex region was separated into individual clusters, something we did not detect using the WT mouse brain nor in the unimputed Mouse AD data (Fig. 4b). Further cortex-analysis revealed a clear separation of the cortex region into two spatially separable parts (Fig. 4c). The first cluster consisted of the cortical subplate, olfactory, entorhinal, ectorhinal, temporal association, and perirhinal areas. The second cluster contained the auditory, primary somatosensory, posterior parietal association, and retrosplenial areas. When these two clusters were mapped to the anatomical reference, the cluster 1 occupied the upper quadrant while cluster 2 occupied the lower quadrant (Fig. 4e, f).

Since such heterogeneity was only detected in the imputed AD cortex but not the imputed WT cortex, we further investigated these two clusters’ roles in AD progression. Specifically, we extracted significantly AD-activated genes for each cluster, respectively, using the Wilcoxon rank-sum test. By selecting upregulated genes in the AD sample with fold change greater than 50% and FDR < 0.01, we identified 55 markers for cluster 1 and 41 for cluster 2 (Ext. Fig. 4.5). We found 34 genes that are only significantly upregulated in AD cluster 1 region (Fig. 4e) and 20 genes that are only significantly upregulated in AD cluster 2 region (Fig. 4f). Only 21 AD activated genes are shared for these two clusters, suggesting the heterogeneity of these two clusters in AD progression (Fig. 4d).

### MIST recovers spatial gene-gene interactions

Spatial gene-gene co-expression plays an important role in understanding gene interactions across 2D space. The dropouts within the ST data undermine the correlation analysis’s power and cause inaccurate estimation of gene-gene spatial correlation. Based on the reference Allen Brain Atlas (ABA) regional gene-expression values, we obtained two pairs of spatially highly correlated genes: *Cldn11-Arhgef10* and *Gfap-Aqp4*^23^. In the ABA data sets, gene *Cldn11* and *Arhgef10* have a spatial PCC of 0.99 and SCC of 0.97 (Fig. 5c). Gene *Gfap* and *Aqp4* have a spatial PCC of 0.72 and SCC of 0.74 (Fig. 5f).

**Figure 5.**
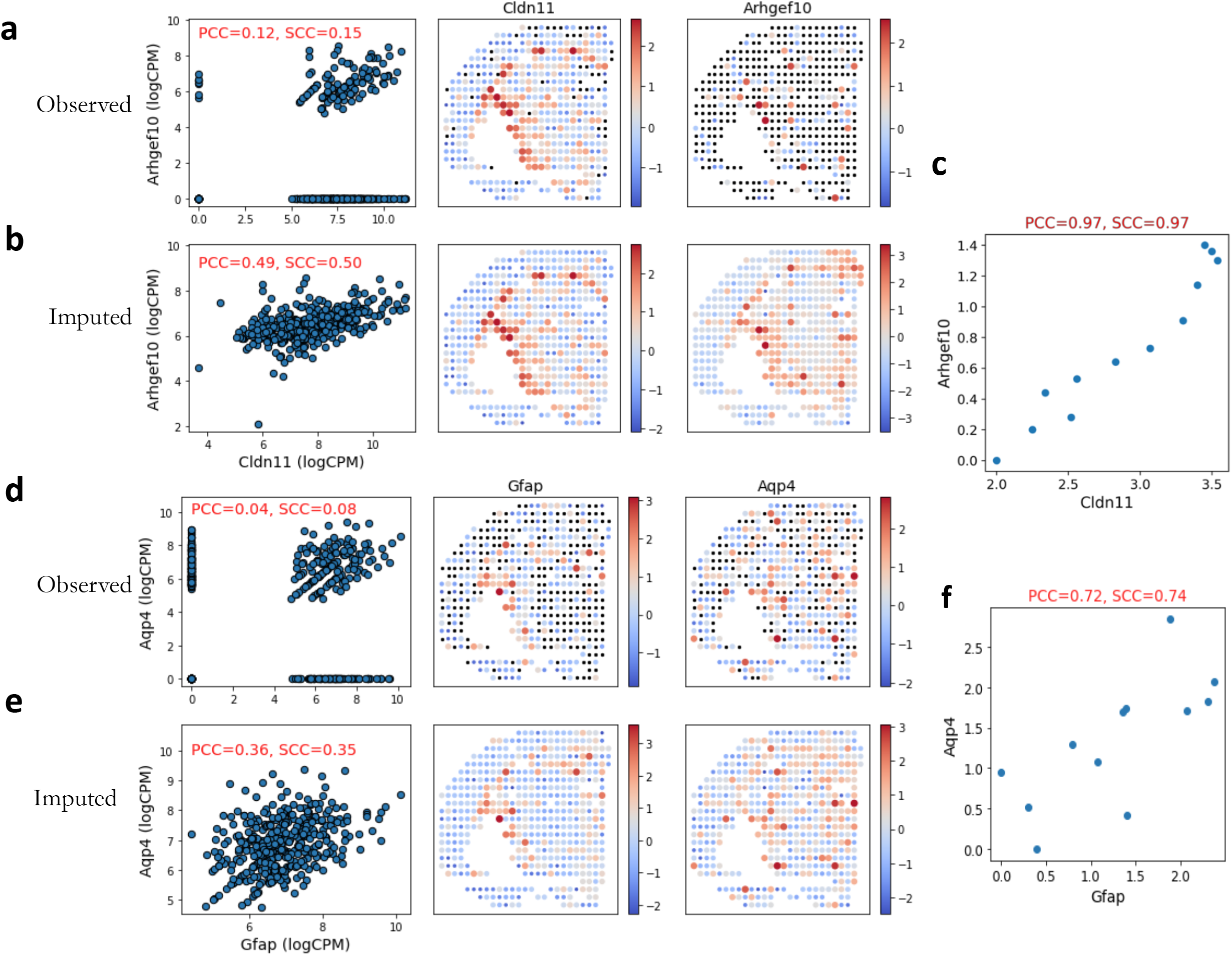
MIST recovers spatially co-expressed gene pairs. **a,** correlation of *Cldn11-Arhgef10* using original observed data. Left, scatter plot of expression values for gene pair regardless of positions. Right, spatial expression pattern of individual genes. **b,** correlation of *Cldn11-Arhgef10* using MIST-imputed data. **c**, correlation of *Cldn11-Arhgef10* using Allen Brain Atlas (ABA) regional gene expression data. **d,** correlation of *Gfap-Aqp4* using original observed data. **e,** correlation of *Gfap-Aqp4* using MIST-imputed data. **f**, correlation of *Gfap-Aqp4* using Allen Brain Atlas (ABA) regional gene expression data.

Using these two pairs of highly spatially correlated genes as examples, we showed that MIST-imputed ST data can recover the lost spatial co-expression signals by accurately estimating the dropout values (Fig. 5). While the non-zero spots’ expression values are visually associated for gene pairs *Cldn11-Arhgef10* and *Gfap-Aqp4*, excessive dropouts reduced the statistical models’ efficacy in estimating the correlations. Before imputation, Cldn11-Arhgef10 only have a Spearman’s correlation coefficient (SCC) of 0.15 with a p-value of 0.002 (Fig. 5a). After imputation, the SCC score was improved to 0.50 with a p-value of 3.5E-29 (Fig. 5b). Moreover, *Gfap-Aqp4* has insignificant correlation (SCC=0.08, p-value=0.56) before imputation. MIST significantly rescued *Gfap-Aqp4*’s correlation with an SCC of 0.31 (p-value=9E-12, Fig. 5d, e).

## Conclusions

We developed a spatial transcriptomics imputation method– MIST–by estimating the missing values for every automatically detected region within the tissue sample. We demonstrated MIST’s efficacy by showing its superior performance at estimating hold-out values across multiple datasets when compared to other imputation methods.

MIST’s ability to accurately recover lost gene-expression does not rely on complicated models. Compared with complicated models like deepImpute^24^, which uses a deep neural network to estimate missing values, MIST assumes that the gene-expression matrix has a low rank. When a relatively small number of cell types reside in a tissue, MIST’s assumption is simple yet interpretable. Compared with scRNA-seq imputation methods such as McImpute that also use a low rank assumption on a data set, MIST utilizes the spatial information that was critical in evaluating the affinity among tissue spots but not accessible in scRNA-seq data.

Though primarily developed to impute the missing values in ST data, MIST addresses another ST-data-analysis challenge: *in silico* region identification. MIST first automatically and accurately detects the tissue regions within the ST data, bypassing the manual annotation process, which can be troublesome, especially when tissue heterogeneity is not easily detectable by human eyes.

We showed that imputing the missing values in ST using MIST is essential to ST data analysis. First, MIST allows discovery of the true spatial co-expressed gene pairs, which are crucial for analyzing spatial gene interaction and cell communication. Second, MIST enhances the heterogeneity within tissue structures and helps detect spatially differentially expressed genes that contribute to such spatial heterogeneity.

We therefore unequivocally recommend using MIST prior to ST downstream analyses, such as when identifying spatial gene-gene interactions. While the original ST data provided by 10X Visium might hinder ST analysis’ accuracy, MIST will accurately rescue the missing values and drastically increase the signal-to-noise ratio. Moreover, our *sui generis in-silico* region detection enables analyzing the ST at a brand-new level that combines anatomical connectivity and molecular similarity. It will be useful in many areas, including identifying local subregions within tumors whose heterogeneity is hard to see through pathological staining images.

## Methods

### Data and preprocessing

The ST data sets we used in this study include samples from a 12-month wild-type mouse brain (MouseWT), a 12-month Alzheimer's Disease mouse brain (MouseAD), a Melanoma tumor, and a Prostate tumor sample^5,6,8^. We first filtered low-quality genes with raw counts <3 in fewer than 2 spots within each sample. Performing this quality-control measure allowed us to remove genes that might otherwise introduce noise to the pipeline. We used a raw mRNA count matrix to measure the gene-expression abundance. But because every spot contains tens of cells, the uncertain cell number and sequencing depth would influence the mRNA abundance detected. We therefore used counter per million (CPM) to normalize the raw mRNA count, where 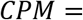 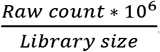.

### Detecting tissue boundaries by identifying local connected neighborhoods (LCNs)

Suppose the ST dataset has *N* genes and *M* spots, the spatial gene-expression profile can be defined as 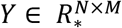, where *Y* is the observed CPM count matrix, and R_*_ denotes the set of all N by M matrices with non-negative real values. The *M* spots in a ST slide can form a lattice graph, *G =<V, E>*, where V is the node-set and *E* is the edge-set. Every pair of adjacent (*u*, *v*) spots are connected with *E (u, v)* representing the weight value.

We used Principal Component Analysis (PCA)^15^ to reduce the *Y* matrix’s dimension to infer the edges’ weights. Specifically, we assigned the edge weight between spot u and v by calculating the Pearson Correlation Coefficient (PCC) between the two spots using the first k principal components that explain ≥ 80% of the total variance. The PCC scores are calculated using Python *SciPy*^25^ package.

Next, we filtered out edges with weights less than ε to remove functionally dissimilar edges. Doing this enabled us to detect tissue boundaries, which are essential to performing accurate imputation using spatial information. We used a depth-first search algorithm^26^ to identify all the connected components in graph *G*. Each local connection component is predicted as an independent tissue region.

### Inferring missing gene-expression values using the average of multiple runs of the rank minimization algorithm

#### Algorithm 1

Ensemble rank minimization

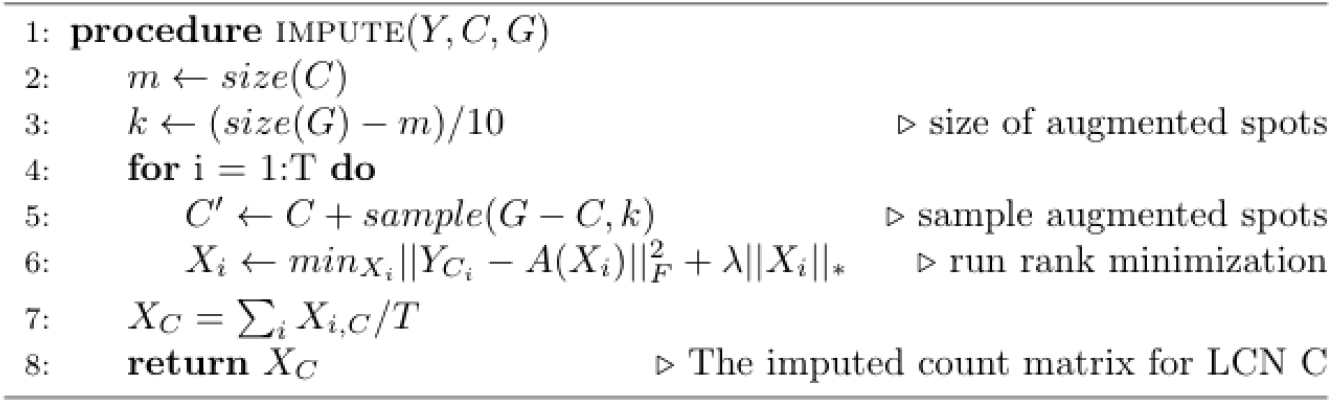

To impute each LCN *C*, we averaged multiple rank minimization algorithm runs on an augmented neighborhood *C* (Algorithm 1). We randomly selected 10% of spots from *G* that are not members in *C* to augment the community *C’* for imputation. In our pipeline, we adopted the convex relaxation of the rank minimization algorithm that was originally implemented in McImpute^12^ to impute the noise-added community *C’*. By repeating this procedure *T* times, we introduced diversity to the community and stabilized it by averaging the imputed values for *C*.

### Random hold-out of highly expressed genes for parameter learning

Using a high threshold while filtering will result in many isolated spots, which will undermine the rank minimization algorithm’s power. On the other hand, using a low threshold will fail to detect tissue boundaries accurately. Therefore, choosing an appropriate threshold for each dataset is critical to the model’s performance.

To automatically select the optimal *ε* for every dataset, we identified a set of high-density genes with non-zero expression values across at least 80% of all spots. We define this as a landmark gene set to optimize the parameter ∊ for our imputation model. Specifically, we randomly withheld 50% of the landmark genes' expression values and observed our model’s imputation accuracy using different *ε* values. We did a grid search for *ε* from 0 to 1 and selected the *ε* with the lowest Rooted Mean Squared Error (RMSE) defined as 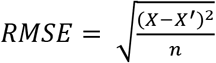, where *X* is the hold-out non-zero values, *X’* represents the MIST estimated values using certain *ε* and *n* denotes the number of non-zero hold-out values.

### Data generation for hold-out experiments

We preprocessed and normalized the MouseWT, MouseAD, Melanoma tumor, and Prostate tumor samples. We then filtered out genes with non-zero proportions less than 50% to guarantee the quality of the remaining genes after further random hold-out.

To generate the hold-out data, we used a 5-fold cross validation schema for the non-zero values. Specially, we first randomly partitioned every gene’s nonzero values into 5 groups. In each hold out, we created missing values by setting one group to zero and performed imputation based on the remaining values. The held-out values served as ground truth for evaluating the imputation algorithms’ accuracy. We reported two metrics, RMSE and PCC, where RMSE measures the estimation error and PCC measures the concordance between the true expression values and the estimated values.

**Table 1.** Data summary for hold-out experiments

|  | *MouseWT* | *MouseAD* | *Melanoma* | *Prostate* |
| --- | --- | --- | --- | --- |
| *#genes (preprocessed)* | 10178 | 11241 | 4498 | 8073 |
| *#genes (<50% sparsity)* | 3024 | 4081 | 952 | 1718 |
| *#spots* | 447 | 488 | 293 | 406 |

## Source Code Availability

The MIST algorithm is implemented in Python and is available at https://github.com/linhuawang/MIST.git.

All the source code and raw data for this manuscript can be downloaded from https://github.com/LiuzLab/MIST_manuscript.git.

## References

1. Marx, V. Method of the Year: spatially resolved transcriptomics. Nat. Methods 18, 9–14 (2021).

2. Asp, M., Bergenstråhle, J. & Lundeberg, J. Spatially resolved transcriptomes—next generation tools for tissue exploration. BioEssays 42, 1900221 (2020).

3. Ståhl, P. L. et al. Visualization and analysis of gene expression in tissue sections by spatial transcriptomics. Science (80-.). 353, 78–82 (2016).

4. He, B. et al. Integrating spatial gene expression and breast tumour morphology via deep learning. Nat. Biomed. Eng. 1–8 (2020).

5. Berglund, E. et al. Spatial maps of prostate cancer transcriptomes reveal an unexplored landscape of heterogeneity. Nat. Commun. 9, 1–13 (2018).

6. Thrane, K., Eriksson, H., Maaskola, J., Hansson, J. & Lundeberg, J. Spatially resolved transcriptomics enables dissection of genetic heterogeneity in stage III cutaneous malignant melanoma. Cancer Res. 78, 5970–5979 (2018).

7. Maynard, K. R. et al. Transcriptome-scale spatial gene expression in the human dorsolateral prefrontal cortex. Nat. Neurosci. 1–12 (2021).

8. Chen, W.-T. et al. Spatial transcriptomics and in situ sequencing to study Alzheimer’s disease. Cell 182, 976–991 (2020).

9. van Dijk, D. et al. MAGIC: A diffusion-based imputation method reveals gene-gene interactions in single-cell RNA-sequencing data. BioRxiv 111591 (2017).

10. Jeong, H. & Liu, Z. PRIME: a probabilistic imputation method to reduce dropout effects in single-cell RNA sequencing. Bioinformatics 36, 4021–4029 (2020).

11. Wagner, F., Yan, Y. & Yanai, I. K-nearest neighbor smoothing for high-throughput single-cell RNA-Seq data. BioRxiv 217737 (2017).

12. Mongia, A., Sengupta, D. & Majumdar, A. McImpute: matrix completion based imputation for single cell RNA-seq data. Front. Genet. 10, 9 (2019).

13. Huang, M. et al. SAVER: gene expression recovery for single-cell RNA sequencing. Nat. Methods 15, 539–542 (2018).

14. Linderman, G. C., Zhao, J. & Kluger, Y. Zero-preserving imputation of scRNA-seq data using low-rank approximation. bioRxiv 397588 (2018).

15. Wold, S., Esbensen, K. & Geladi, P. Principal component analysis. Chemom. Intell. Lab. Syst. 2, 37–52 (1987).

16. Chen, Y.-T. et al. Serological analysis of Melan-A (MART-1), a melanocyte-specific protein homogeneously expressed in human melanomas. Proc. Natl. Acad. Sci. 93, 5915–5919 (1996).

17. Xie, S. et al. Expression of MCAM/MUC18 by human melanoma cells leads to increased tumor growth and metastasis. Cancer Res. 57, 2295–2303 (1997).

18. Zhou, Y. et al. Osteopontin expression correlates with melanoma invasion. J. Invest. Dermatol. 124, 1044–1052 (2005).

19. Shomali, N. et al. Heat shock proteins regulating toll-like receptors and the immune system could be a novel therapeutic target for melanoma. Curr. Mol. Med. 21, 15–24 (2021).

20. Yu, G., Wang, L.-G., Han, Y. & He, Q.-Y. clusterProfiler: an R package for comparing biological themes among gene clusters. Omi. a J. Integr. Biol. 16, 284–287 (2012).

21. Hou, W., Ji, Z., Ji, H. & Hicks, S. C. A systematic evaluation of single-cell RNA-sequencing imputation methods. Genome Biol. 21, 1–30 (2020).

22. McInnes, L., Healy, J. & Melville, J. Umap: Uniform manifold approximation and projection for dimension reduction. arXiv Prepr. arXiv1802.03426 (2018).

23. Jones, A. R., Overly, C. C. & Sunkin, S. M. The Allen brain atlas: 5 years and beyond. Nat. Rev. Neurosci. 10, 821–828 (2009).

24. Arisdakessian, C., Poirion, O., Yunits, B., Zhu, X. & Garmire, L. X. DeepImpute: an accurate, fast, and scalable deep neural network method to impute single-cell RNA-seq data. Genome Biol. 20, 1–14 (2019).

25. Virtanen, P. et al. {SciPy} 1.0: Fundamental Algorithms for Scientific Computing in Python. Nat. Methods 17, 261–272 (2020).

26. Tarjan, R. Depth-first search and linear graph algorithms. SIAM J. Comput. 1, 146–160 (1972).

