## Extended Figures for "Missing-value imputation and *in-silico* region detection for spatially resolved transcriptomics"

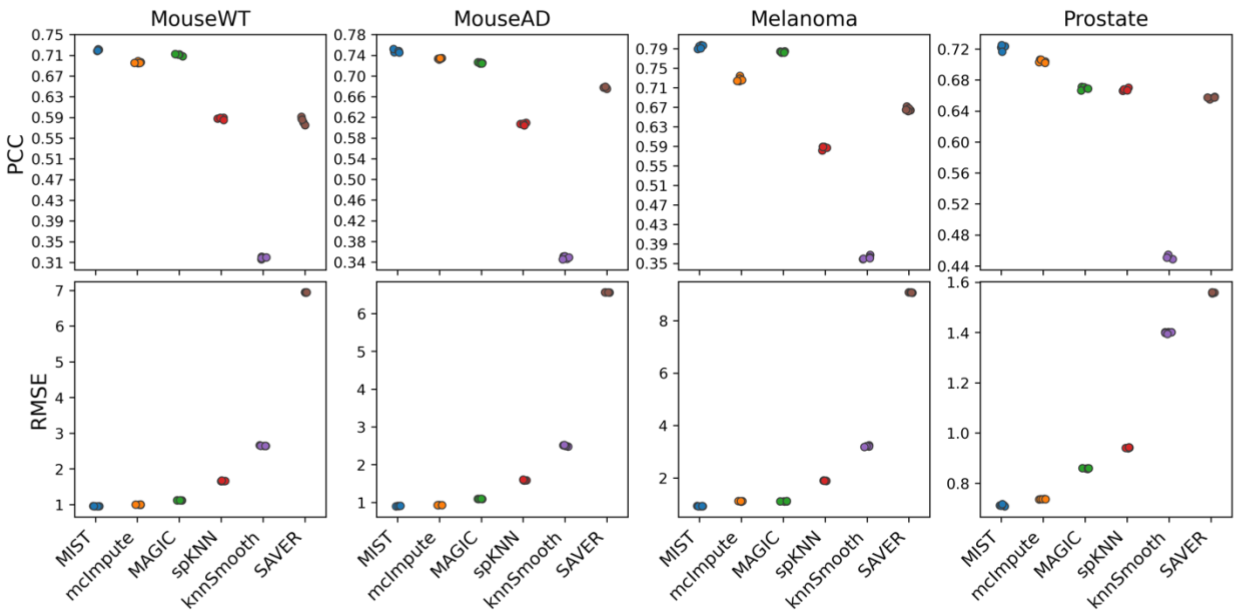


**Extended Figure 3.1 | Holdout test performance. a-b,** Holdout experiment performance across multiple data sets using metrics **a,** PCC and **b,** RMSE. Each column is an individual data set. Points in the same color indicate the 5 non-overlap fold of tests for each model.

**a**

**b**

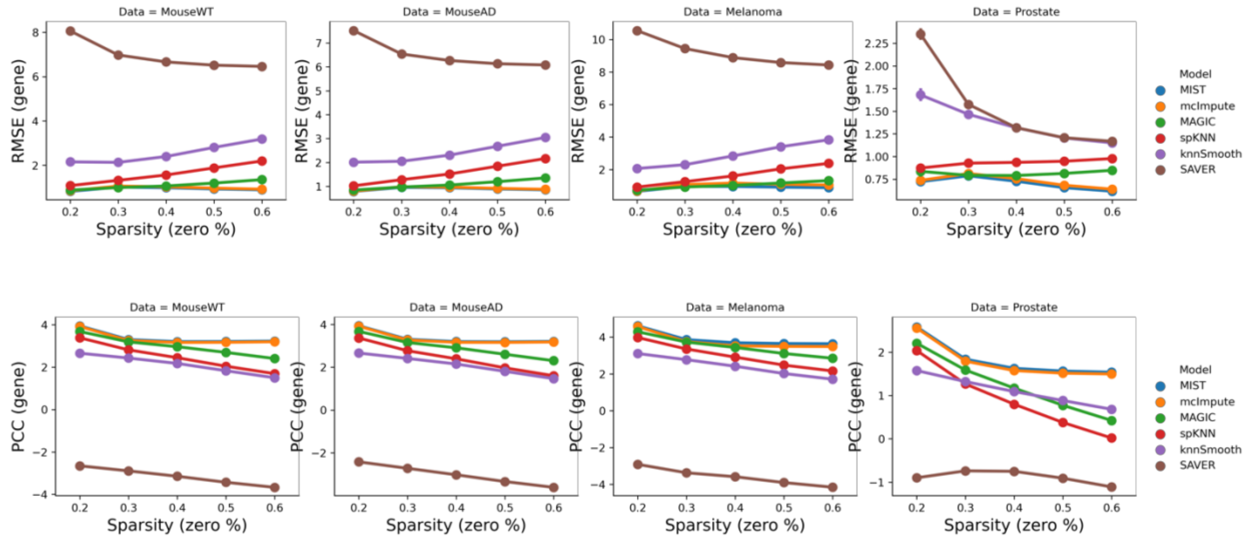


**Extended Figure 3.2 | Gene-level holdout test performance. a.** RMSE (y-axis) for every gene as a function of gene sparsity (x-axis). b. PCC for every gene as a function of gene sparsity.

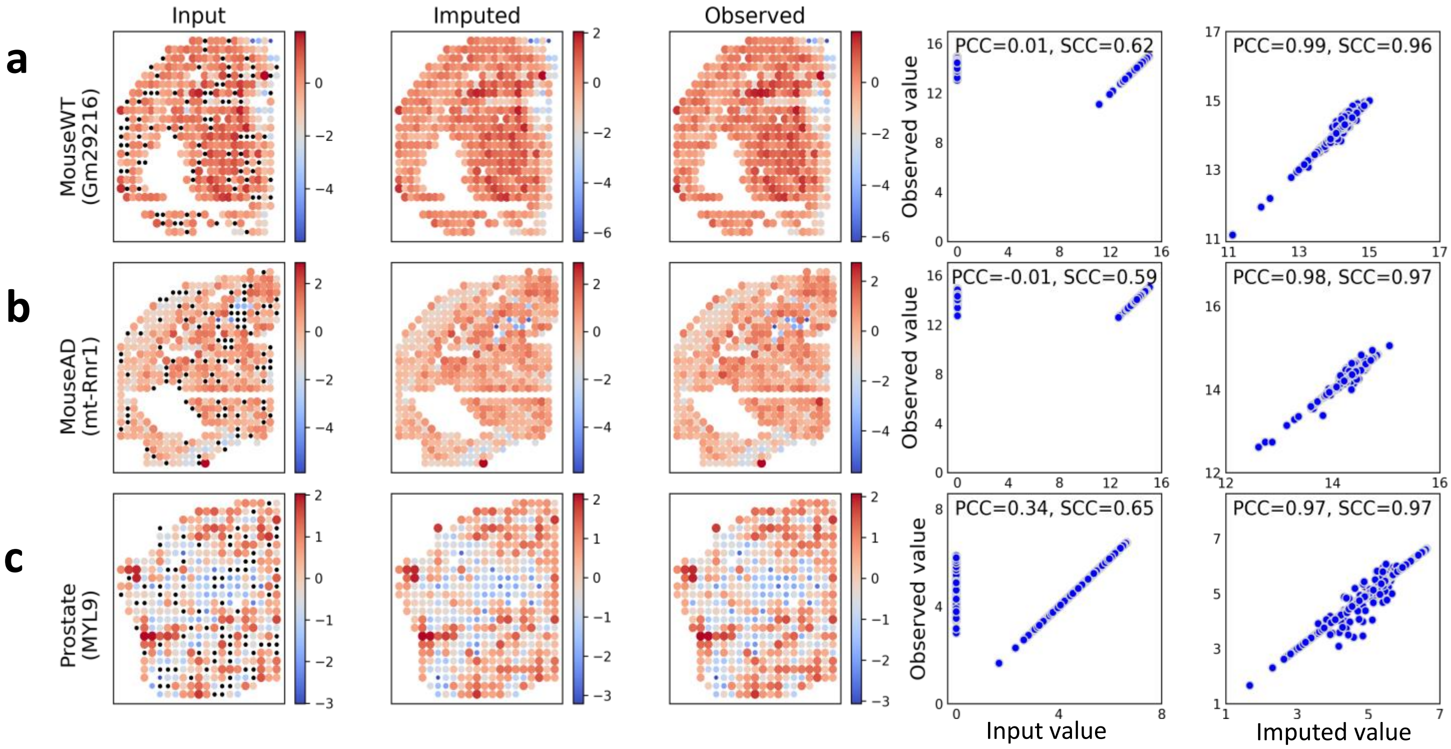


**Extended Figure 3.3 | MIST recovered gene expression patterns in holdout experiments.** Column 1 is the spatial pattern of the input to MIST with random non-zero values held out. Column 2 is the imputed gene expression pattern. Column 3 is the original observed gene expression pattern. Column 4 shows the correlation between input values and observed values. Column 5 shows the correlation between the imputed expression values and the observed expression values. **a,** Expression patterns recovered for gene *Gm29216* in the Mouse WT brain. **b,** Expression patterns recovered for gene mt-Rnr1 in the Mouse AD brain. **c,** Expression patterns recovered for gene *GM29216* in the Prostate sample.

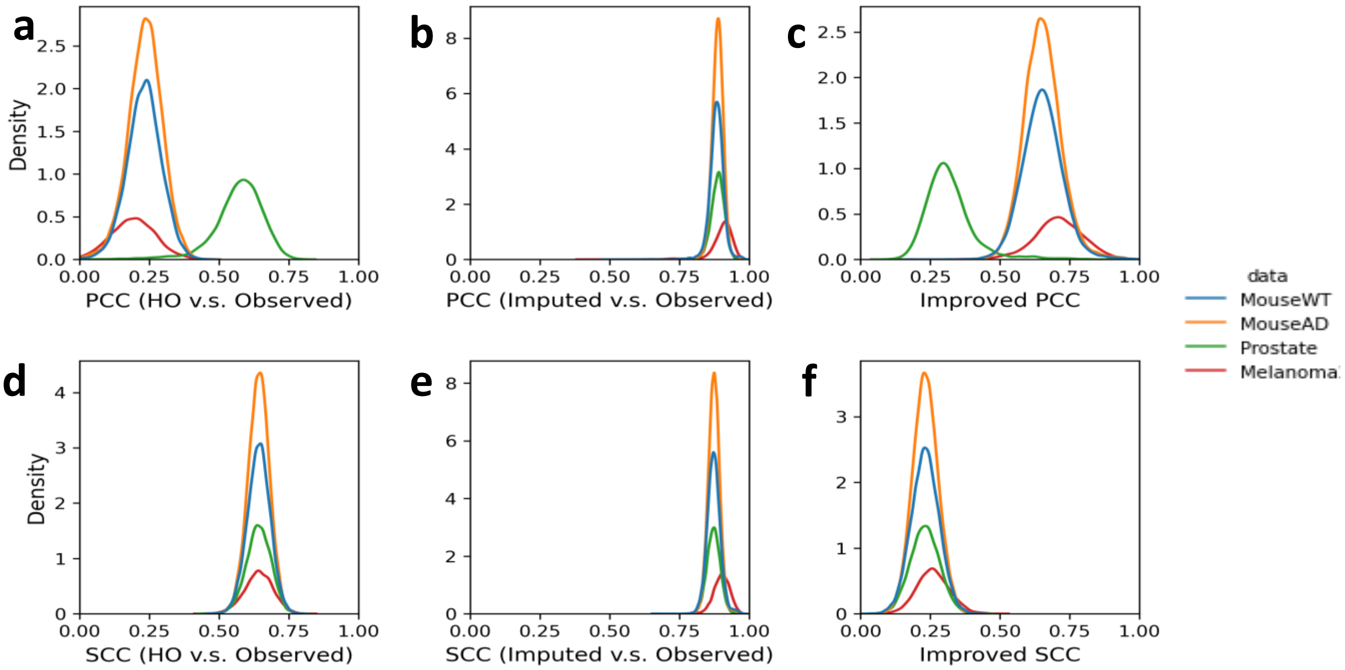


**Extended Figure 3.4 | Distribution of improved PCC and SCC in holdout experiments. a,** PCC distribution of HO vs. Observed non-zero gene expression. **b,** PCC distribution of Imputed vs. Observed non-zero gene expression. **c,** Distribution of improved PCC for nonzero values of each held out gene. **d,** SCC distribution of HO vs. Observed non-zero gene expression. **e,** SCC distribution of Imputed vs. Observed non-zero gene expression. **f,** Distribution of improved SCC for nonzero values of each held out gene.

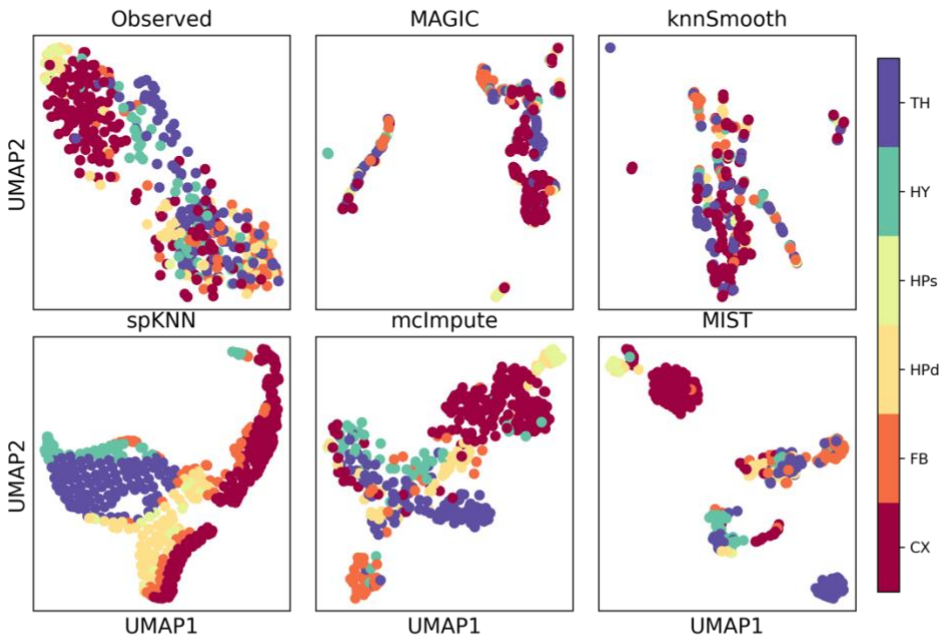


**Extended Figure 4.1 | UMAP visualization of Mouse WT brain using data imputed by different algorithms.**

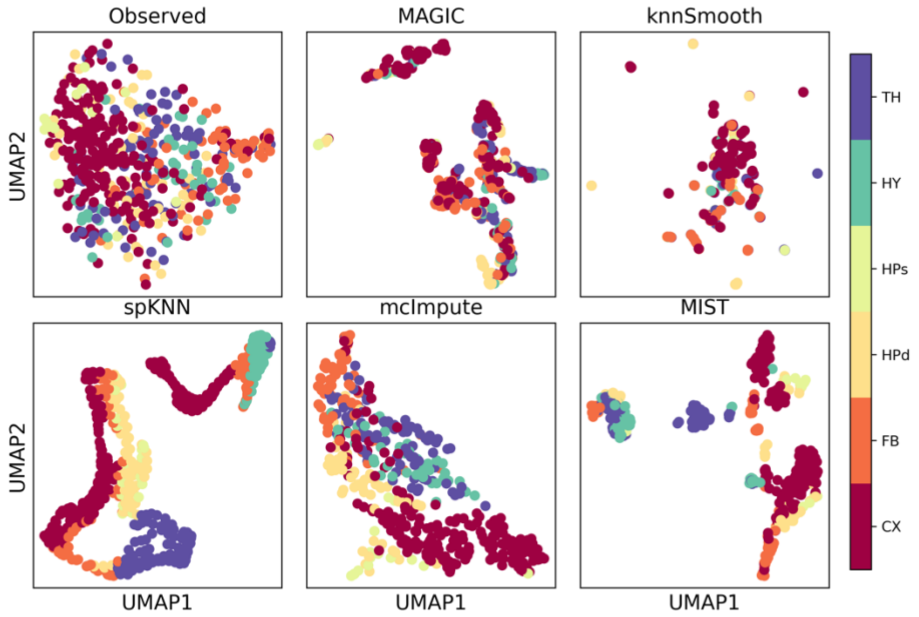


**Extended Figure 4.2 | UMAP visualization of Mouse AD brain using data imputed by different algorithms.**

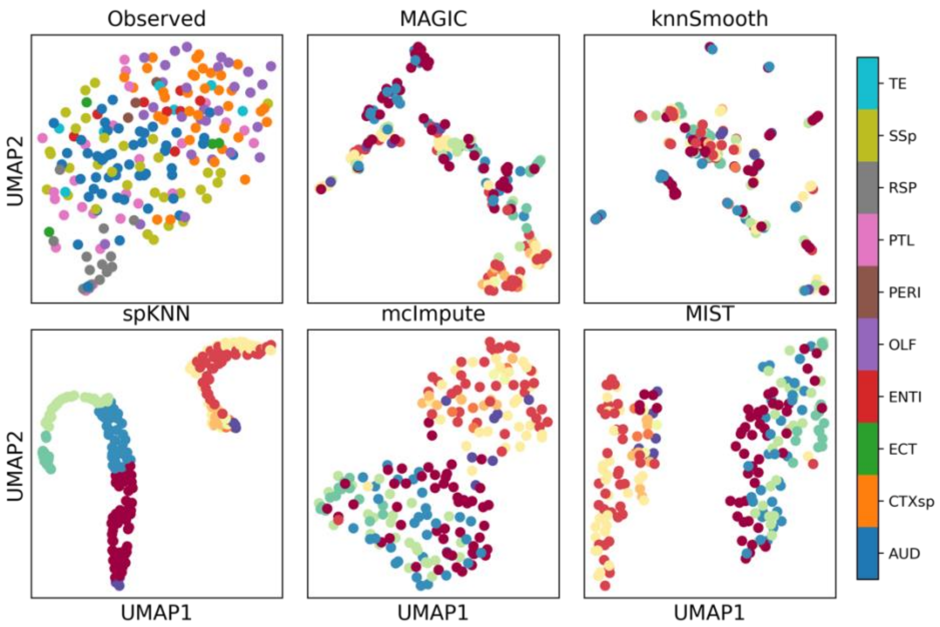


**Extended Figure 4.3 | UMAP visualization of Mouse AD Cortex using data imputed by different algorithms.**

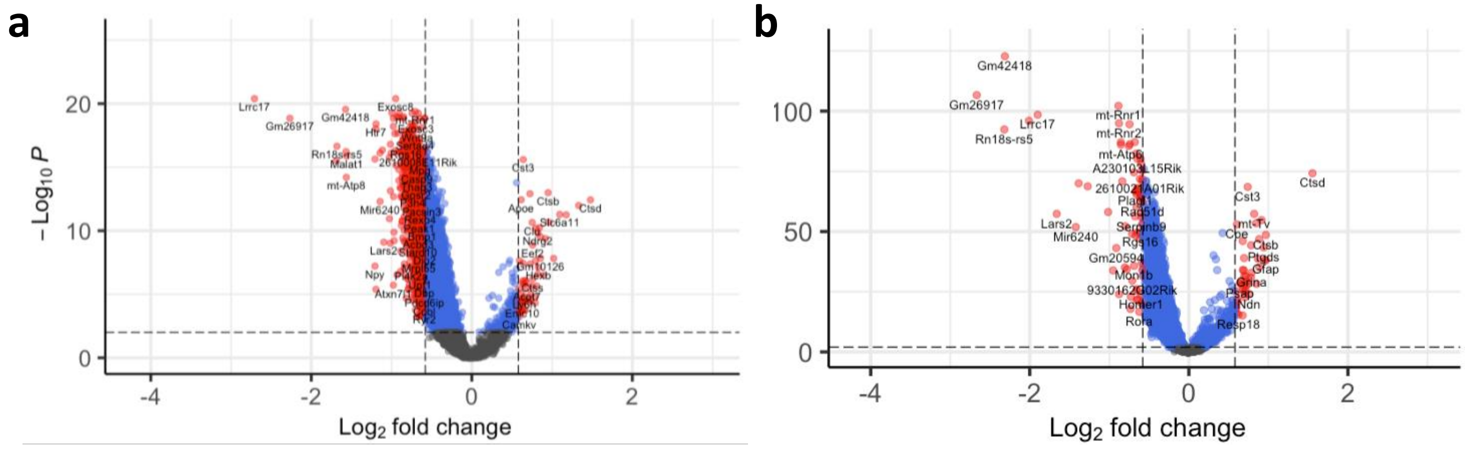


**Extended Figure 4.4 | Volcano plots of AD vs. WT differential expression analysis. a,** Volcano plot of differential gene expression analysis results comparing AD vs. WT mouse brain using Cluster 1 MIST-imputed data. Red dots are significant genes with absolute LFC > 0.58 and FDR < 0.01. **b,** Volcano plot of differential gene expression analysis results comparing AD vs. WT mouse brain using Cluster 2 samples.

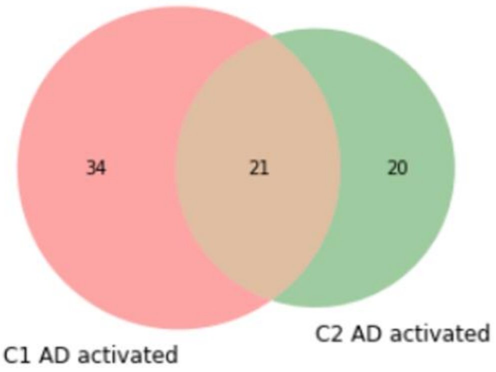


Slc6a11 Ndrg2 Sparc Gm10126 Oaz1 Hexb Ctss 6330403K07Rik Atp6ap1 Lamp1 Tmx4 Tyrobp Wdr6 Acot7 Bex3 Selenom Ly6h Ahi1 Sncb Rpl28-ps3 Cldn11 Hap1 Laptm4a Uqcrc1 Prdx6 Hbb-bs Emc10 Fasn Ddb1 Zmat2 Rpl8 Scg3 Gm15427 Camkv
Cst3 Ctsb mt-Tv Apoe Ctsd Gfap B2m Ndn Clu Gtf2h1 Ptgds Grina Eef2 Resp18 Gaa Psap mt-Tl2 Trf Nnat Bsg Rnf187
Cpe Dnm1 Tuba1a Slc22a17 Ttr Gng3 Enpp2 Dynll2 mt-Th Tcf25 Gabbr1 Rps15 Pebp1 Syngr1 Pld3 RP23-269H21.1 Eno2 Eef1a2 Pip5k1a Sez6

**Extended Figure 4.5 | Consensus and difference of AD activated genes between spatial cortex clusters.** Circle and gene symbols in red color indicate genes that are only activated in Cluster 1 (C1, CTXsp, OLF, ENTI, TE, ECT and PERI). Circle and gene symbols in green are genes that are only activated in Cluster 2 (C2, AUD, PTL, RSP, SSp). Orange color indicates AD activated genes that are shared across these two spatial cortex clusters.
